# Computational Prediction of Multiple Antigen Epitopes

**DOI:** 10.1101/2024.08.08.607232

**Authors:** R. Viswanathan, M. Carroll, A. Roffe, J.E. Fajardo, A. Fiser

**Affiliations:** Department of Chemistry and Biochemistry, Yeshiva College, NY, NY 10033, USA; Department of Chemistry and Biochemistry, Stern College for Women, NY, NY 10016, USA; Department of Systems and Computational Biology, Albert Einstein College of Medicine, Bronx, NY, 10461, USA

## Abstract

**Motivation:** Identifying antigen epitopes is essential in medical applications, such as immunodiagnostic reagent discovery, vaccine design, and drug development. Computational approaches can complement low-throughput, time-consuming, and costly experimental determination of epitopes. Currently available prediction methods, however, have moderate success predicting epitopes, which limits their applicability. Epitope prediction is further complicated by the fact that multiple epitopes may be located on the same antigen and complete experimental data is often unavailable.

**Results:** Here, we introduce the antigen epitope prediction program ISPIPab that combines information from two feature-based methods and a docking-based method. We demonstrate that ISPIPab outperforms each of its individual classifiers as well as other state-of-the-art methods, including those designed specifically for epitope prediction. By combining the prediction algorithm with hierarchical clustering, we show that we can effectively capture epitopes that align with available experimental data while also revealing additional novel targets for future experimental investigations.

**Supplementary information:** Supplementary data are available at *Bioinformatics* online.

## 1 Introduction

Antibodies (or immunoglobulins) are protein molecules that recognize and bind to specific regions of an antigen, known as epitopes (Van Regenmortel 2009). The identification of the epitopes involved in antibody-antigen interactions is particularly important in the field of immunology and essential in medical applications, such as therapeutic antibody development (Melo, Lemos et al. 2018; Garofalo M 2020), immunodiagnostic reagent discovery (Milich 1989; Leinikki, Lehtinen et al. 1993), and vaccine design (Van Regenmortel 2006; Dudek, Perlmutter et al. 2010; Robinson and Mulligan 2016; Yang, Zhang et al. 2016; Palatnik-de-Sousa, Soares et al. 2018). Epitopes are classified as either linear or conformational. A linear epitope is a short peptide of continuous residues in a protein sequence. A conformational epitope, on the other hand, consists of spatially proximal residues in the three-dimensional structure of the folded protein, and it is composed of residues that are at least partially discontinuous in the sequence.

Conformational epitopes constitute approximately 90% of all known cases (Ansari and Raghava 2010). Conformational epitopes cannot be simulated by a short peptide of amino acid sequences as they only exist embedded in the entire protein structure, and its binding activity cannot be measured outside of the protein context. Consequently, structural analysis of antibody-antigen complexes must be performed to successfully map these epitopes.

Several experimental techniques are employed to map epitopes from antibody-antigen complexes, including NMR spectroscopy (O’Connell, Gamsjaeger et al. 2009), X-ray crystallography (Kobe, Guncar et al. 2008; Shi 2014), and cryo-EM (Callaway 2015). Each method has its own unique advantages and weaknesses, but overall, experimental determination of epitopes is low-throughput, time-consuming, costly and often experimentally inaccessible (Zheng, Liang et al. 2023).

Computational methods have emerged as an alternative option to complement experimental techniques to identify protein interfaces. While many of these methods are more general, some have been developed to specifically predict antigen epitopes (Kringelum, Lundegaard et al. 2012)El-Manzalawy, Dobbs et al. 2008; Tomar and De 2014; Sanchez-Trincado, Gomez-Perosanz et al. 2017). As discussed by Zheng et al. (Zheng, Liang et al. 2023), computational methods to predict protein-protein interfaces are generally classified as either sequence or structure-based approaches. Sequence-based methods consider several properties of the query protein beyond its primary structure, including its flexibility, secondary structure, hydrophilicity, solvent accessibility, and antigenicity. These methods begin by using a sliding window between 3-30 residues wide, with the target residues situated at the midpoint. Many currently available methods rely on a combination of physicochemical properties for predicting linear epitopes like BcePred (Saha, Bhasin et al. 2005), BEPITOPE (Odorico and Pellequer 2003), and PEOPLE (Alix 1999). These methods differ in the features employed and the weighting scales of the properties included over the sliding window used. The next generation of methods use machine learning algorithms and statistical approaches, including random forest and support vector machines, which are trained and optimized with experimentally determined linear epitopes listed in B-cell epitopes databases, such as Bcipep (Saha, Bhasin et al. 2005), and IEDB (Peters, Sidney et al. 2005). Most of the currently available epitope prediction methods fall into this category and thus are primarily focused on identifying linear epitopes. Other methods use SVM classifiers (Chen, Liu et al. 2007), neural network model, ABCpred (Saha and Raghava 2006), or Hidden Markov Model (Larsen, Lund et al. 2006) to predict linear epitopes. A recent evaluation of the available machine learning algorithms found that SVMTriP performed the best out of the available methods with respect to specificity and accuracy (Galanis, Nastou et al. 2021).

Structure-based approaches perform epitope predictions in the context of the three-dimensional structure of an antigen. These methods are especially important because more than 90% of identified epitopes are conformational. Furthermore, protein structures are dynamic, and their native conformations can depend on whether they are complexed or uncomplexed with an antibody (Brown, Joaquim et al. 2011). Zheng et al. suggested that structural methods are superior to sequence-based methods as they can consider these possibilities. However, far fewer prediction methods exist for discontinuous epitopes due to their greater design complexity (Zheng, Liang et al. 2023). Within structure-based approaches, two sub-classes exist. The first sub-class of methods are the “template-free” approaches that incorporate both sequential features— including hydrophobicity, physicochemical properties, amino acid types and evolutionary conservation information — as well as structural features — including geometric shape, solvent-accessible surface area, and secondary structure — of a query protein (Gallet, Charloteaux et al. 2000; Ofran and Rost 2003; Yan, Dobbs et al. 2004; Esmaielbeiki, Krawczyk et al. 2016). These methods are trained and optimized on experimentally determined protein structures to balance sequential and structural features with the probability that a given residue is interfacial. The second sub-class of methods are the “template-based” approaches that map the structure of a query antigen onto homologous structures, whose interfaces have previously been determined, to predict interfacial residues (Zhang, Deng et al. 2011). However, the development of reliable structural methods is limited by the availability and quality of both the three-dimensional structure of the query and its homologs with known interfaces (Esmaielbeiki, Krawczyk et al. 2016). Some of the currently available epitope-specific models include CEP(Kulkarni-Kale, Bhosle et al. 2005), DiscoTope 2.0 (Haste Andersen, Nielsen et al. 2006), DiscoTope 3.0 (Hoie, Gade et al. 2024), ElliPro (Ponomarenko, Bui et al. 2008), EPCES (Liang, Zheng et al. 2009), EpiPred (Krawczyk, Liu et al. 2014), EPITOPIA (Rubinstein, Mayrose et al. 2009; Liang, Zheng et al. 2010), EPSVR (Liang, Zheng et al. 2010), PEASE (Sela-Culang, Ashkenazi et al. 2015), PEPITO (Sweredoski and Baldi 2008), SEPIa(Dalkas and Rooman 2017), and SEPPA(Sun, Wu et al. 2009). Independent analyses have found that these structural models perform quite similarly to one another but have low overall accuracy in identifying antigen epitopes (Zhang, Niu et al. 2012; Yao, Zheng et al. 2013).

Protein-protein interface prediction approaches can be further classified as “partner-specific” or “partner-independent.” Partner-specific approaches refer to methods that require the structures or sequences of both interacting proteins, while other methods can predict interfacial residues on a query protein alone without the knowledge of its cognate partner and are thus partner-independent. Since the identity or knowledge of the cognate antibody is not always known in practice, all methods discussed in this work are partner-independent. Meta prediction methods are useful approaches that integrate two or more individual computational classification models and output a consensus interface prediction. Meta-methods differ in the specific orthogonal information that is combined and in the way it is optimized. For example, meta-PPISP uses linear regression analysis to combine cons-PPISP (Chen and Zhou 2005), PROMATE (Neuvirth, Raz et al. 2004), and PINUP (Liang, Zhang et al. 2006; Esmaielbeiki, Krawczyk et al. 2016). These individual classifiers are all template-free approaches. A more recent method, VORFFIP, a complex structure-based method, integrates heterogeneous data, such as residue level structural and energetic features, evolutionary sequence conservation, and crystallographic B-factor using random forest approach (Segura, Jones et al. 2011).

We recently reported the development of an integrated method for protein interface prediction, ISPIP (Walder, Edelstein et al. 2022). We found that the efficacy of an integrated method could be improved by using a suitable combination of individual classifiers that rely on orthogonal structure-based properties of query proteins. ISPIP integrates ISPRED4 (Savojardo, Fariselli et al. 2017), PredUs 2.0 (Zhang, Deng et al. 2011), and DockPred (Viswanathan, Fajardo et al. 2019), which are template-free, templated-based, and docking-based partner-independent predictors, respectively, through simple linear or logistic regression or more advanced random-forest machine learning algorithms, such as XGBoost. On a diverse test set of 156 query proteins, ISPIP outperformed each of the models that it integrates as well as other state-of-the-art methods in identifying protein interfaces. In this work, we present a new method, ISPIPab, to predict antigen epitopes based on the framework of ISPIP.

In computational epitope prediction, complications arise owing to the possibility of multiple epitopal regions on a single antigen. Therefore, it is prudent to analyze the predicted epitope residues to identify potentially distinct epitope regions that can be investigated experimentally. Previous work has attempted to cluster predicted residues into potential epitopes following computational prediction. Zhang et al. (Zhang, Zhao et al. 2014) describes the development of CBEP, a method, which combines sequence features and selects the optimal subset of features from the high dimensional feature space using the Fisher-Markov selector. CBEP uses machine learning classification algorithms with an added cost value that penalizes the wrong identification of classes. Additionally, a K-means clustering algorithm is used to group the antigenic residues into clusters based on their spatial location and the value of the threshold parameter. Ren et al. presents a staged heterogeneity learning algorithm that uses only sequential information and various patterns of propensities for epitope prediction, as well as a clustering method with a pre-defined optimal cutoff distance of 6 Å between any pair of residues belonging to the same cluster, as this was found to yield the highest F-scores (Ren, Song et al. 2017).

Our current study integrates three individual classifiers (SPPIDER, ISPRED4, and DockPred) using XGBoost (Breiman 2001; Pedregosa 2011), an optimized random forest algorithm, to predict antigen epitopes. Furthermore, in order to identify all possible putative epitopal regions on an antigen, we implemented a hierarchical clustering algorithm in ISPIPab, with an optimal number of clusters determined based on the predicted epitopal residues for each antigen. We found that the current method ISPIPab (Integrated structure-based protein interface prediction for antibody binding) outperforms each of its individual classifiers, SPPIDER, ISPRED4, and DockPred as well as VORFFIP, meta-PPISP, SEPPA 3.0, and DiscoTope 2.0 in predicting antigen epitopes.

## 2 Methods

### 2.1 Databases

#### 2.1.1 Bound antigen dataset A

This dataset is from Jespersen et al. and consists of 335 antibody-antigen complexes with experimentally identified epitope residues (Jespersen, Mahajan et al. 2019). This set includes the three-dimensional structures from the IEDB database (Vita, Overton et al. 2015) and unpublished complexes in the Protein Data Bank (PDB) (Berman, Westbrook et al. 2000) that were found using antibody-specific Lyra Hidden Markov Models. These complexes include only those with B-cell heavy-light chain receptors, antigens of greater than 60 residues, and structures with a resolution of <3 Å. Redundancies were removed by clustering the antibodies and antigens at 90% and 70% sequence identity thresholds, respectively. This resulted in 202 antibody-antigen clusters. We further refined this dataset by removing redundancy at a stricter level and retained only complexes in which the antigens shared ≤30% sequence identity. This reduced the number of complexed antigens to 107. We refer to this dataset as “Bound dataset A.”

#### 2.1.2 Unbound antigen dataset B

Complexed antibody-antigen structures require knowledge of a cognate antibody, which is often unavailable from experimental studies. Besides, there may be conformational changes during antibody-antigen complex formation that are not discernable in the uncomplexed antigen. Therefore, to be more general, we sought to test our method using only the three-dimensional structure of the unbound antigen.

To accomplish this, we searched for uncomplexed antigen structures analogous to those in the bound complexes. We identified uncomplexed antigen structures in the PDB with >95% sequence identity to the antigens in the bound structures (in dataset A) and composed of only one distinct protein entity. We identified 76 uncomplexed monomer antigens while ensuring a sequence identity threshold of ≤30%. We refer to this dataset as “Unbound dataset B.” Datasets A and B, composed of bound and unbound antigens, respectively, were used to assess the performance of ISPIPab on both types of antigen structures.

To further expand our unbound dataset, we searched for structures in the SACS database (Allcorn and Martin 2002) that contained an antibody-antigen complex, a resolution of <3 Å, antigens between 100-450 residues, and were published after 2005. This returned 2196 structures. The epitopes in these complexes were determined using CSU program (Sobolev, Sorokine et al. 1999) with a cutoff value of 4.0Å between any atom in a residue in the antigen and an atom in the antibody of the complex and establishes a legitimate contact type according to CSU. After applying identical constraints to those in the unbound dataset B, we identified 35 additional unbound antigens, providing a total of 111 antigens for our study. This expanded dataset (dataset B2) of unbound antigens includes the original dataset B and the additional 35 antigens. The epitope residues in dataset B were identified from dataset A by sequentially aligning them.

Dataset B2 consists of proteins of varying lengths (Fig. S1) and are representative of different protein families (Fig. S2). 59 antigens are included in Fig. S2. An additional 35 antigens each belong to a unique, different topology and hence are grouped by their architecture (Fig. S3). 19 of the 111 antigens do not have an assigned CATH classification (Orengo, Michie et al. 1997).

#### 2.2.1 ISPIPab and Individual Classifiers

ISPIPab is a meta-method that integrates the predictions of three independent classifiers (SPPIDER, ISPRED4, and DockPred) to predict epitope residues on query antigens.

SPPIDER uses a novel approach by incorporating enhanced relative solvent accessibility (RSA) predictions, utilizing the difference between predicted and observed RSA to identify interaction sites. The work demonstrated that RSA-based fingerprints surpass other features like conservation, physicochemical properties, and structural data. While SPPIDER is a structure-based interface prediction method and not template-based, it leverages complexes from several homologous proteins to determine the composite interfacial region for the query protein. SPPIDER integrates a total of 19 features, including sequential, evolutionary, structural, and RSA-based factors, into its model using a combination of machine learning techniques (SVM, NN, and LDA) for optimal interface prediction (Porollo and Meller 2007).

ISPRED4 is a template-free interface predictor that uses a SVM model trained on a dataset of 314 distinct monomer chains whose complex structures were resolved by X-ray crystallography. Accordingly, residues whose accessible surface area decrease by at least 1 Å^2^ when comparing their unbound and complex states are considered interfacial. Interface classification is accomplished using a 46-dimensional feature vector consisting of 10 groups of descriptors, including 34 sequence-based and 12 structure-based features, with both sets each forming five descriptor groups (Savojardo, Fariselli et al. 2017).

DockPred (Viswanathan, Fajardo et al. 2019), a docking-based model, was developed based on the hypothesis that proteins, like small organic molecules, tend to bind to energetically favorable sites on a protein, regardless of their biological cognate partners. This model simulates the docking of a query protein on 13 non-cognate proteins that are distinct with respect to size and fold, and the results demonstrated that non-cognate protein ligands similarly bind to cognate binding sites of target proteins.

ISPIPab (Fig. 1.) employs XGBoost, a machine learning algorithm utilizing an ensemble of decision trees and gradient boosting, to integrate normalized interface likelihood scores from three or more individual classifiers for each residue, generating consensus epitope predictions. Cutoff values for each classifier and tree level are optimized, and model parameters, such as maximum tree depth and pruning parameter α, are fine-tuned for optimal fit.

**Fig. 1.**
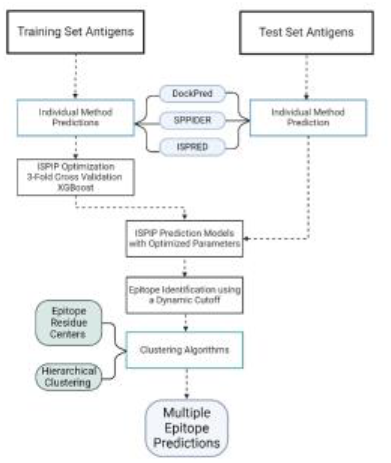
Schematics of ISPIPab to determine multiple epitopes on antigens.

### 2.3 Interface Prediction

The prediction methods used in this study return interface likelihood scores (p) that range between 0 and 1 for each residue. To determine the threshold for the number of top scoring residues considered as epitopal in each antigen, we used a dynamic threshold. As proposed by Zhang et al., a dynamic threshold, N, can be calculated using the following equation: *N*=6.1 *R*^0.3^, where R is the number of surface-exposed residues on the antigen (Zhang, Deng et al. 2011).

In Fig. S4, we compare the total number of epitope residues determined either experimentally or using CSU with the dynamic threshold for each antigen. The dynamic cutoff is significantly higher than the total number of annotated residues in all cases. But since we do not have prior knowledge of the number of epitope residues for any antigen of interest, using the dynamic threshold to obtain a set of predicted epitope residues from the computational method is necessary.

### 2.4 Performance Evaluation

Once the computationally predicted epitopes are known, the elements of the confusion matrix, True Positive (TP), True Negative (TN), False Positive (FP) and False Negative (FN) can be determined by comparison with the experimentally known epitopal residues.

To assess the predictive performance of the individual classifiers and other recent methods in addition to ISPIPab, we used the F1-score (F1), Matthew’s correlation coefficient (MCC), and the areas under the receiver operating characteristic (ROC-AUC) and precision-recall (PR-AUC) curves. To assess whether the F1-scores were normally distributed and test the null hypothesis, the nonparametric Kolmogorov–Smirnov (KS) single- and two-sample tests were used. The differences in F1 scores and MCC were considered statistically significant if the p-value of the KS test was <0.05.

### 2.5 Training and Testing Sets

The antigens in unbound dataset B2 were partitioned into three training sets and a test set. The test set was composed of 29 antigens while the three training sets were each composed of 27-28 antigens. The training and test sets consisted of a similar distribution of antigen sizes and CATH families. The size distributions in the training and test sets are shown in Fig. S5. Through three-fold cross-validation, the predictions of SPIDDER, ISPRED4, and DockPred were integrated using XGBoost.

### 2.6 Determining the number of independent epitopes by hierarchical clustering

The computationally predicted set of N epitope residues were clustered using hierarchical clustering based on the distances between their geometric centers. Using the Scikit-Learn library, agglomerative hierarchical clustering was performed (Pedregosa 2011). The linkage criterion was set to ‘ward’ so as to minimize the variance of the clusters that were merged, and the Euclidean distance was the set metric to calculate pairwise distances between any two residues’ geometric centers with no preset distance threshold.

The optimal number of clusters for each antigen corresponds to the number of vertical lines in the dendrogram cut by a horizontal line that can traverse the maximum distance vertically without intersecting a cluster. This method allows us to determine the optimal number of clusters dynamically. As seen in Fig. 2, the optimal number of clusters for the antigen with PDB ID 3F5V is 3. The horizontal line drawn at around 31 cuts three vertical lines (corresponding to # clusters =3) and can traverse to 71 vertically, traversing a vertical distance of 40 without intersecting a cluster. The horizontal line at 71 (corresponding to # clusters =2) can only traverse vertically without intersecting a cluster to 88, a vertical distance of 17. Thus, the optimal number of clusters in this case is determined to be 3.

**Fig. 2.**
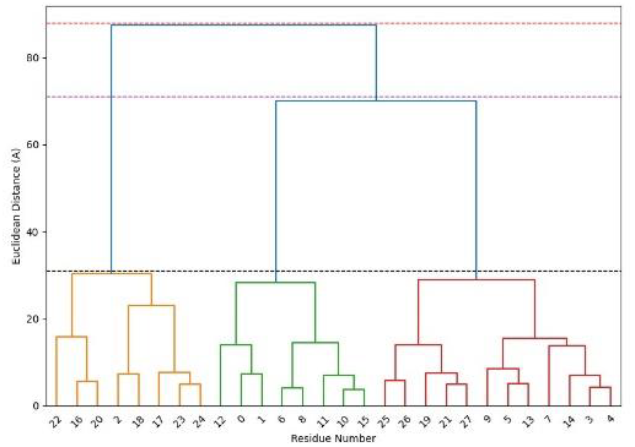
Dendrogram for PDB ID 3F5V showing the dynamic selection of the number of clusters.

## 3 Results and Discussion

We explored the possibility of identifying multiple epitopes on an antigen without knowing its cognate antibody partner by predicting epitopal residues on a set of antigens with ISPIPab followed by hierarchical clustering, with the number of clusters determined dynamically for each antigen. We could successfully identify multiple epitopes through a single calculation on an antigen using its three-dimensional structure. This is illustrated in Fig. 3 for HIV-1 envelope GP120, which has two experimentally determined complexes with two different antibodies that show two distinct epitopal regions. The total number of predicted interfacial residues (N=32) predicted by ISPIPab are clustered into two binding regions (indicated by red and green spheres in Fig. 3) and these capture the two distinct experimentally identified epitopes very well.

**Fig. 3.**
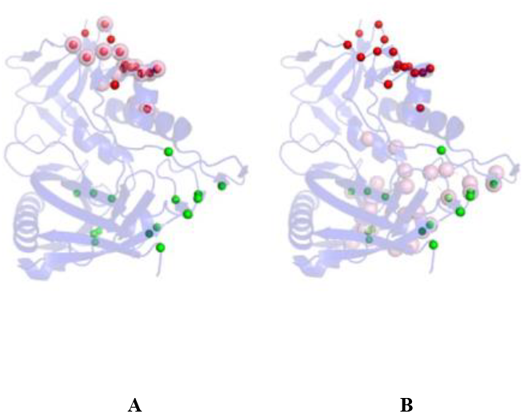
HIV-1 GP120 with the two experimentally identified epitopes (in pink). Panel A shows the antigen complex with ADCC-potent antibody N60-i3 Fab (PDB ID: 5KJR). while panel B shows the antigen complex with a broadly neutralizing antibody (PDB ID: 4JPW) The epitopal residues predicted by ISPIPab are clustered into Cluster 1 (red) and Cluster 2 (green) and compared with the experimentally identified epitopes (pink).

Cluster 1, as indicated by the red spheres in Fig. 3A (PDB ID: 5KJR), accurately captures the binding site for HIV-1 GP120 with ADCC-potent antibody N60-i3 Fab (pink spheres Fig. 3A) with an F1-score of 0.79 while cluster 2, as indicated by the green spheres in Fig. 3B, (PDB ID: 4JPW), accurately captures the binding site for HIV-1 GP120 with the broadly neutralizing antibody bNAb (pink spheres Fig. 3B) with an F1-score of 0.48.

### 3.1 Comparison between Bound and Unbound Antigens

To determine if there were any significant differences in the performance of epitope prediction using the bound versus unbound antigen structures, the analogous datasets A and B of bound and unbound antigens described earlier were evenly distributed amongst the training and testing sets for three-fold cross validation. Each of the testing sets consisted of 16 antigens while the training sets for Bound dataset A were slightly larger than those for Unbound dataset B. The performance of ISPIPab was assessed on these sets using the F1-score and MCC metrics (Table 1). ISPIPab performs equally well on Unbound dataset B when trained with either the bound or unbound datasets. Overall, the predictions on Unbound dataset B have a slightly higher F1-score and MCC compared to the predictions on Bound dataset A.

**Table 1.** Comparison of average F1-score and MCC for an analogous test set of bound and unbound antigens using bound or unbound antigen training sets.

| Training Set | Bound dataset A | Bound dataset A | Unbound dataset B |
| --- | --- | --- | --- |
| Test Set | Bound dataset A | Unbound dataset B | Unbound dataset B |
| < F1-score > | 0.299 | 0.350 | 0.335 |
| <MCC> | 0.214 | 0.283 | 0.264 |

The comparable performance of ISPIPab on a test set of unbound antigens using a training set of bound or unbound antigens shows that the method is successful in predicting epitopes without the structural information of the binding antibody partner. Although conformational changes are expected during antibody-antigen binding, the results in Table 1 suggest that epitope predictions are possible without complete knowledge of these conformational changes.

### 3.2 Performance of Integrated Method versus Individual Classifiers

We explored the performance of the method on Dataset B2 consisting of 111 unbound antigens. The dataset was evenly distributed among 3 training sets (27-28 antigens each) and a testing set of 29 antigens. As shown in the Methods section, the distributions of antigen size (Fig. S5) and CATH family classification were uniform across the training and test sets. The performance of ISPIPab is significantly better than any of the individual classifiers that were integrated (Table 2.). The average F1-score of ISPIPab is 0.312, while the average F1-scores of the individual classifiers range from 0.122 to 0.194. The F1-scores and MCC do not follow a normal distribution as tested by a single-sample Kolmogorov-Smirnov (KS) test. The statistical significance of the differences in F1-scores and MCC between ISPIPab and the other individual classifiers were verified using a two-sample KS test at a 95% confidence level. All the p-values were significantly smaller than 0.05.

**Table 2.** Comparison of average F1-score and MCC of three individual classifiers with ISPIPab. Statistical significance was verified using a two-sample KS test at a 95% confidence level. The p-values range from 0.0001 - 0.0149.

|  | SPPIDER | ISPRED | DockPred | ISPIPab |
| --- | --- | --- | --- | --- |
| <F1 score> | 0.122±0.13 | 0.194±0.18 | 0.168±0.14 | 0.312±0.16 |
| <MCC> | 0.003±0.15 | 0.089±0.20 | 0.054±0.16 | 0.230±0.17 |

As seen in Table 2, the standard deviations are large for all individual classifiers. This indicates large protein-to-protein variations in these methods and that each will fail for some antigens. The integration of these methods by ISPIPab, while still having a large standard deviation, takes advantage of the best performing method in each case.

The performance of ISPIPab was also monitored by the area under the Receiver-Operator Curve (AUC-ROC) and Precision-Recall (AUC-PR) curves. The AUC-ROC for ISPIPab is 0.77 while it ranges from 0.62 to 0.66 for the other individual classifiers. Similarly, the AUC-PR for ISPIPab is 0.23 while it ranges from 0.10 to 0.13 for the other individual classifiers. By all statistical measures used, ISPIPab has a considerably stronger predictive power than each of the three individual classifiers.

### 3.3 Importance of all Three Classifiers in the Integrated Model

We next addressed the question of the importance of each individual classifier in improving the performance of ISPIPab. We evaluated the performance of ISPIPab by integrating only two of the three classifiers into the XGBoost algorithm (Table 3). Each of these individual classifiers considers different structural and sequence features, and the predictive power of the integrated algorithm increases when all three are included. While the average F1-score of the ISPIPab model that integrates all three individual classifiers is 0.312, when one of the three is left out, the average F1-score decreases to a range from 0.244 to 0.263. A similar trend is observed for the MCC values. Although it is clear that integrating all three individual classifiers improves performance, it is not clear if all three classifiers contribute equally to enhance the performance of ISPIPab.

**Table 3.** Average F1-scores and MCC of ISPIPab by integrating all three or any two individual classifiers.

| Individual Classifiers Included | SPPIDER, DockPred and ISPRED | SPPIDER and ISPRED | DockPred and ISPRED | DockPred and SPPIDER |
| --- | --- | --- | --- | --- |
| <F1-score> | 0.312±0.16 | 0.246±0.17 | 0.244±0.17 | 0.263±0.16 |
| <MCC> | 0.230±0.17 | 0.153±0.16 | 0.148±0.17 | 0.172±0.17 |

Similar trends were observed with the ROC and PR curves as ISPIPab performs significantly better when all three individual classifiers are integrated as opposed to when one of them is left out. The AUC-ROC ranges from 0.698 to 0.706 and the AUC-PR ranges from 0.161 to 0.171 for the integration of two classifiers.

To highlight the power of ISPIPab, we show a specific example of a membrane protein TRIC (PDB ID: 5H35) (Fig. 5) where the three individual classifiers each yield a poor F1-score. However, ISPIPab integrates the best predictions of each classifier to yield an improved F1-score of 0.45.

**Fig. 4.**
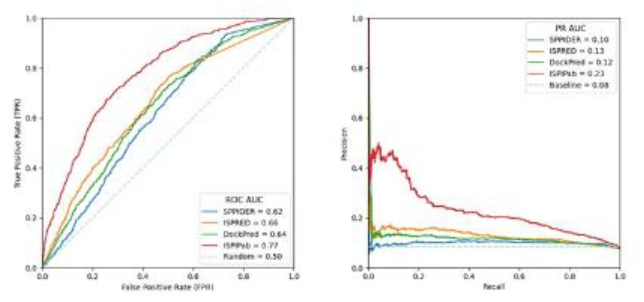
ROC and PR curves comparing the performance of ISPIPab with the individual classifiers.

**Fig. 5.**
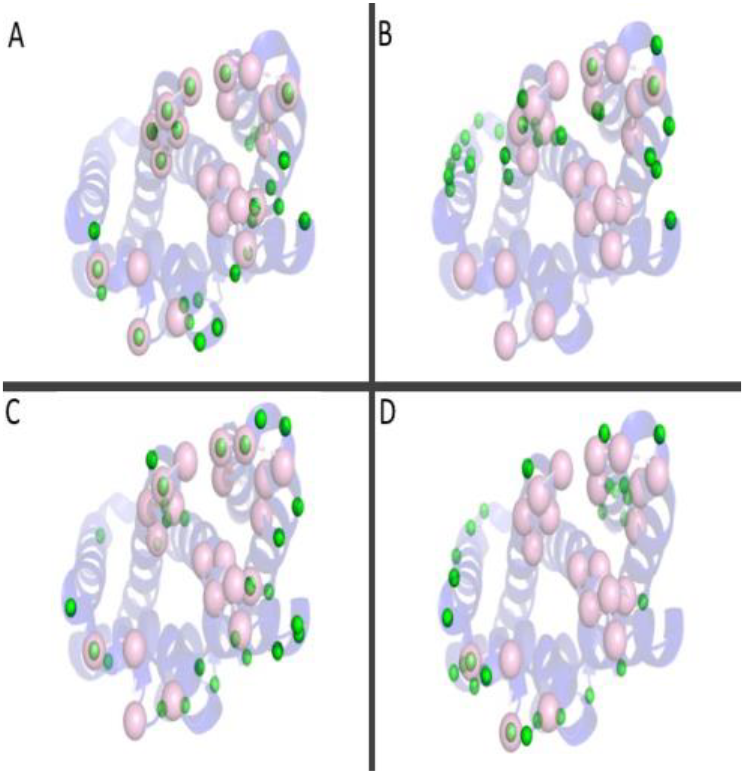
Epitopes predicted by ISPIPab and individual classifiers (green spheres) are compared with the experimental epitope (pink spheres). A. ISPIPab (F1-score = 0.42); B. SPPIDER (F1-score =0.08); C. DockPred (F1-score = 0.04); D. ISPRED (F1-score =0.21).

ISPIPab is also seen to perform better than other state-of-the art methods for predicting interfaces, including VORFFIP (Segura, Marin-Lopez et al. 2015), meta-PPISP (Qin and Zhou 2007), SEPPA 3.0 (Zhou, Chen et al. 2019),DiscoTope 2.0 (Kringelum, Lundegaard et al. 2012) and DiscoTope 3.0 (Hoie, Gade et al. 2024). While VORFFIP and meta-PPISP were developed to identify protein interfaces in general, DiscoTope and SEPPA 3.0 are specifically designed to predict antigen epitopes. DiscoTope 2.0 was shown to be a top scoring method in a recent review (Cia, Pucci et al. 2023) and DiscoTope 3.0 has been shown to perform better than the recent model BepiPred (Clifford, Hoie et al. 2022). The comparison of our results using ISPIPab with DiscoTope is, therefore, meaningful.

The performance of ISPIPab in epitope prediction exceeds those of the other methods, including DiscoTope 2.0 and SEPPA 3.0, with an F1-score of 0.312 and an MCC of 0.230 (Table 4). These differences are statistically significant based on the p-values (<0.05) from a KS two-sample test. Although ISPIPab’s F1-score and MCC are greater than that of DiscoTope 3.0, the difference is not statistically significant at the 95% confidence level according to the KS two-sample test. ISPIPab also has a higher ROC-AUC and PR-AUC than the other methods (Table 4).

**Table 4.**
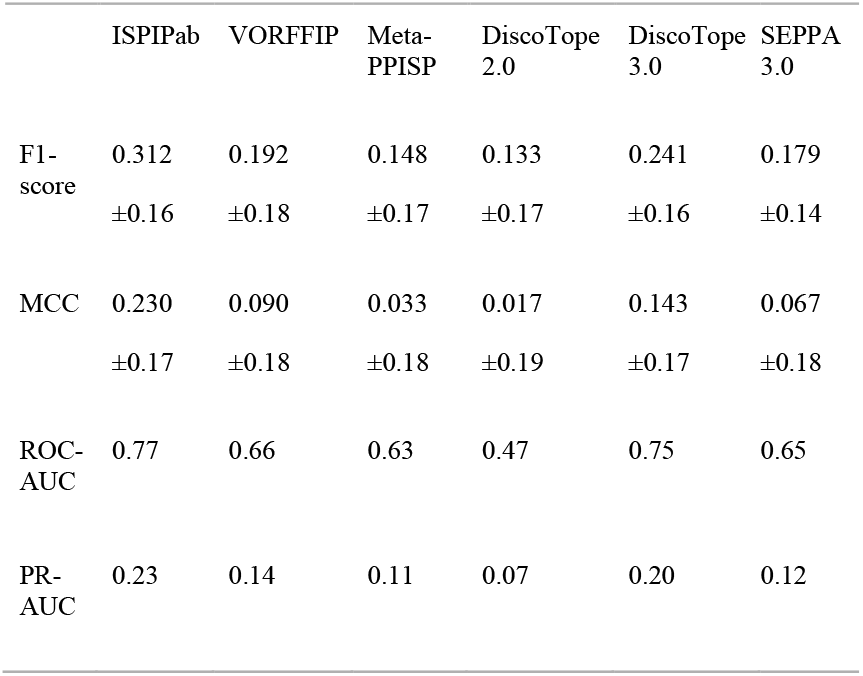
Comparison of average F1-score, MCC, ROC-AUC and PR-AUC of ISPIPab with other methods. The statistical significance was tested using a KS two-sample test, and the p-values range from 0.0027 to 0.0149 for comparison to VORFFIP, meta-PPISP and DiscoTope 2.0 and SEPPA 3.0. For DiscoTope 3.0, the p-value was 0.2926.

|  | ISPIPab | VORFFIP | Meta-PPISP | DiscoTope 2.0 | DiscoTope 3.0 | SEPPA 3.0 |
| --- | --- | --- | --- | --- | --- | --- |
| F1-score | 0.312<br>±0.16 | 0.192<br>±0.18 | 0.148<br>±0.17 | 0.133<br>±0.17 | 0.241<br>±0.16 | 0.179<br>±0.14 |
| MCC | 0.230<br>±0.17 | 0.090<br>±0.18 | 0.033<br>±0.18 | 0.017<br>±0.19 | 0.143<br>±0.17 | 0.067<br>±0.18 |
| ROC-AUC | 0.77 | 0.66 | 0.63 | 0.47 | 0.75 | 0.65 |
| PR-AUC | 0.23 | 0.14 | 0.11 | 0.07 | 0.20 | 0.12 |

### 3.4 Performance Based on Fixed Threshold

The effectiveness of ISPIPab compared to the other methods to identify epitopes is also confirmed based on the number of correctly predicted epitope residues in the top 10, 20, 30, 40, or 50 ranked residues. The results for % TP for the antigens in the test set, calculated using % TP = (Total TP over test set)/(Total annotated residues for test set)*100 are compared in Fig. 6. By this metric as well, ISPIPab performs considerably better than the other methods. This demonstrates that the comparison of the different methods whether based on the dynamic threshold or a fixed threshold will lead to similar conclusions.

**Fig. 6.**
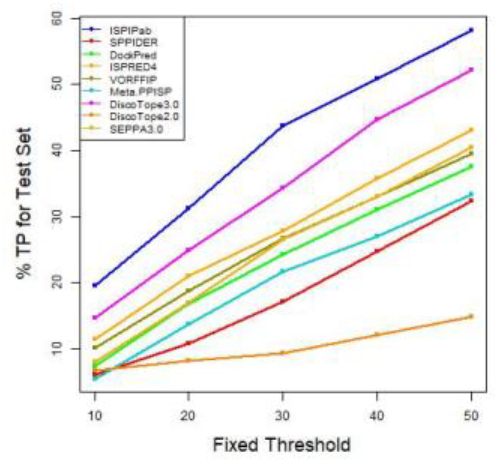
Performance of Methods Based on a Fixed Threshold.

### 3.5 Clustering Improves Prediction Performance of Single Epitopes

We hypothesized that the performance of ISPIPab could be improved by spatial clustering of the predicted epitopes. This was motivated in part by the observation that the dynamic threshold for the interfacial residues, N, is always larger than the number of experimentally determined epitopal residues (Fig. S4). Therefore, the predicted set of epitopal residues could span several spatial locations across the antigen and could predict putative epitopes that have not yet been determined experimentally. We determined the optimal number of spatial clusters for each antigen using hierarchical clustering. One of these clusters of residues often matches well with the experimental epitopes from a particular antibody-antigen complex. The other cluster(s) is a predicted epitope based on the calculations for which experimental data are not yet available. Verification of the other predicted epitope(s) would require additional experiments with complexes of the antigen with various antibodies.

For the testing set of 29 antigens in Dataset B2, hierarchical clustering determined the optimal number of clusters to be 2 for 26 antigens and 3 for the remaining 3 antigens. The distribution of the interfacial residues, N, amongst these clusters is shown in Fig. 7. The number of residues predicted in each epitopal region ranges between 6 and 28 and compares well with the previously reported number of amino acid residues in epitopes, which range from 6 to 29, with an average of 15 and a standard deviation of 4 (Kringelum, Nielsen et al. 2013).

**Fig. 7.**
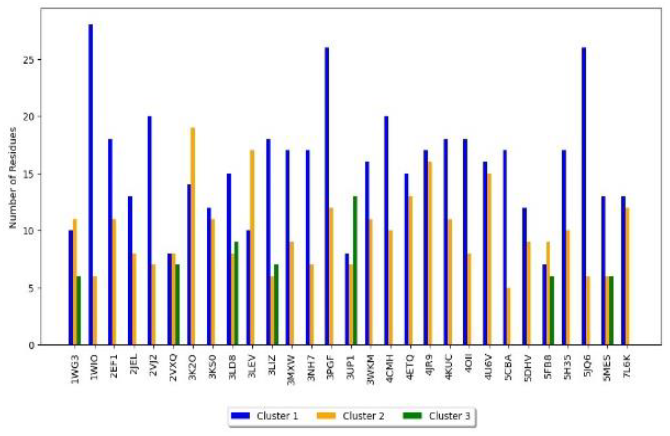
Number of residues in each cluster based on the optimal number of clusters determined by hierarchical clustering. The F1-scores improved upon clustering in all cases. This indicates that including spatially separated residues as part of the “true positive” class when experimental results are only available for a single antibody-antigen complex yields lower scores. Fig. 8 compares the F1-scores of the 29 testing set antigens before and after clustering. In all but 3 cases, the F1-scores improved upon clustering and in a few other cases the F1-score improved from <0.5 to between 0.6 and 0.8. In the three cases where the F1-score decreased, the centers of the two clusters were close enough to be considered a single spatially spread-out epitope.

**Fig. 8.**
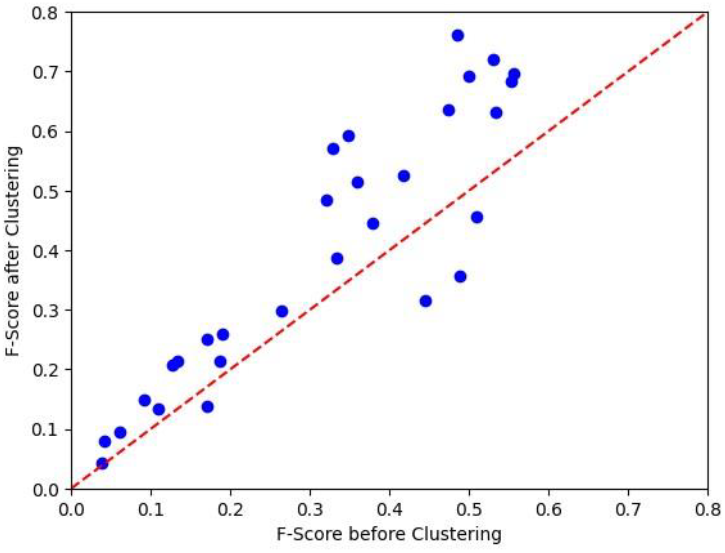
Comparison of F1-scores for the 29 antigens with and without hierarchical clustering. The red line corresponds to no change in F1-score upon clustering.

### 3.6 Hierarchical Clustering Predicts Multiple Epitopes

To illustrate the effectiveness of hierarchical clustering, we further explored the performance of ISPIPab followed by clustering on a set of antigens with multiple experimentally identified epitopes. Starting with an additional set of 31 antigens that were experimentally complexed with different antibodies, 86 complexes were identified. Upon analyzing the epitopes identified in these different complexes, we found that multiple complexes with the same antigen from different experiments had identical epitopes. Therefore, we used a subset of these 31 antigens that did not have identical epitopes. This resulted in a set of 14 antigens with 41 experimentally reported complexes with different antibodies. Upon further analysis of the epitopes of these complexes, it was found that many of the epitopes still had significant overlap.

Complexes of the same antigen have different degrees of residue overlap, as calculated by the percentage of identical residues between the different complexes. The antigens shown in Fig. 9 (PDB ID: 1ALU, 4KXI, 3TGT, and 1IK0) have residue overlaps ranging from 0% to 62%.

**Fig. 9.**
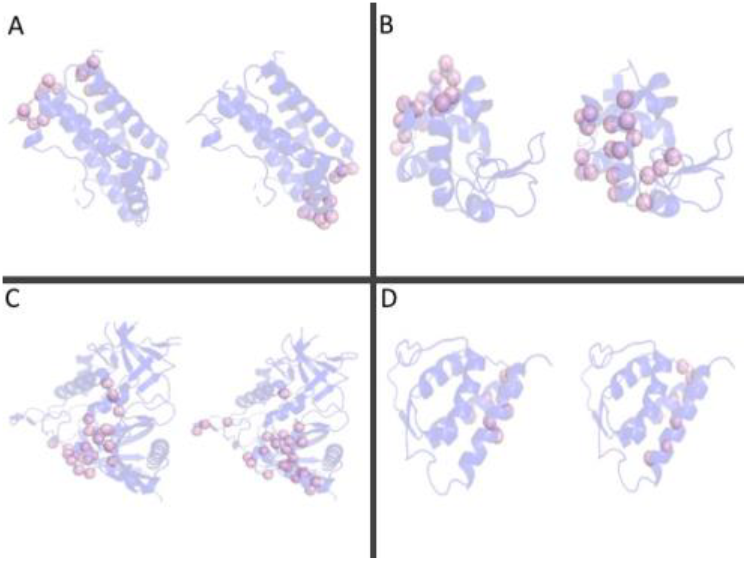
Experimental epitope residues for complexes of antigens with different antibodies with different degrees of overlaps between the epitopes are shown. The pink spheres are the experimentally determined epitopes. A. PDB ID 1ALU, 0% overlap; B. PDB ID 4KXI, 6% overlap; C. PDB ID 3TGT, 25% overlap; D. PDB ID 1IK0, 62% overlap.

The multiple epitopes identified in complexes with different antibodies are in the same spatial region even for epitopes with just 6% overlap. Therefore, we used a 0% residue overlap as the cutoff to be considered as spatially distinct epitopes.

This resulted in 14 antigens with a total of 30 different complexes. The distribution of F1-scores from ISPIPab for each of these 30 complexes is shown in Fig. 10.

**Fig. 10.**
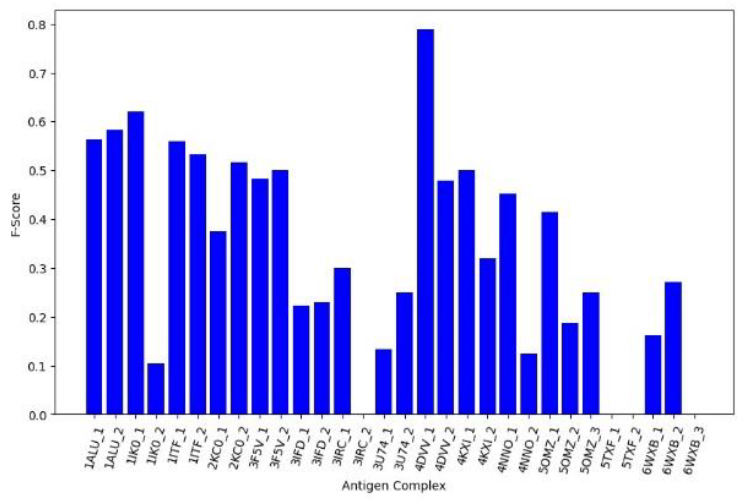
Distribution of F1-scores for the antigens with multiple epitopes.

The average F1-score for these 30 complexes with non-overlapping epitopes is 0.33. It should be noted that the low average F1-score is because of the failure of the method to identify the epitopes in some of the complexes, as seen in both complexes of 5TXF, as well as in one complex from each 3IRC and 6WXB.

### 3.7 Lysozyme as Case Study for Antigens with Multiple Epitopes

Lysozyme is an interesting example with six different complexes with different antibodies and the experimental structures of these complexes are available. For the unbound lysozyme (PDB ID: 4KX1), the dynamic threshold is calculated to be 24. Five of these complexes (PDB ID: 1BVK, 1C08, 2EIZ, 1A2Y, and 4TSA) have epitope residues that overlap between 7% and 88%, and the sixth complex (PDB ID: 1MLC) is distinct with no overlap to the other five.

For the unique complex (1MLC), the total number of experimentally annotated residues is 14. ISPIPab identifies the top 24 scoring residues as the epitope and these residues are clustered using a cluster size of 2, cluster1 and cluster2, with 11 and 13 residues, respectively One of these clusters has an F1 score of 0.32, while the other has an F1 score of 0. Four of the 14 annotated epitopes are identified in cluster1.

For the five complexes where the experimental epitopes overlap, the total number of unique annotated residues identified in these five complexes is 40. Some of these residues occur in multiple complexes. Among the residues that occur in three or more complexes, 6/8 residues are accurately identified by ISPIPab. Of the residues that occur in only one or two of the five complexes, 11/32 residues are accurately identified. Of the 11 residues in cluster1, 7 are TP, and 10 out of 13 residues in cluster2 are TP. Clusters 1 and 2 have F1-scores of 0.27 and 0.38, respectively. Considering that the dynamic threshold (24) is much less than the total number of annotated residues (40), these F1-scores are very good.

ISPIPab with hierarchical clustering certainly captures the multiple likely regions for antibody biding, even in a notorious case like lysozyme. Fig.11 shows the epitopes captured by cluster1 (green) and cluster2 (red) and compares them with the annotated residues (pink). The sizes of the annotated residues are scaled to represent the number of complexes in which it is observed. Fig. 12 shows cluster1 (green) along with the annotated residues in the unique complex. In this unique complex, the antibody is binding to the loop region of the lysozyme.

**Fig. 11.**
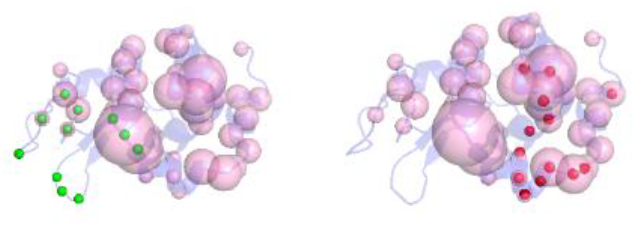
Predicted epitopes in cluster1 (green) and cluster2 (red) and the annotated residues (pink) of the five complexes. The sizes of annotated residues are scaled to the number of complexes where they occur.

**Fig. 12.**
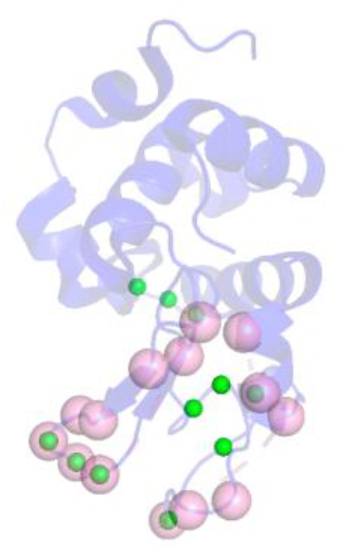
Predicted (green) and annotated residues (pink) in the unique complex, 1MLC. The four TP residues are in the loop region of the antigen.

In general, ISPIPab followed by clustering successfully identifies multiple epitopes. In the cases where the method failed, two of the antigens, 5TXF and 6WXB, are larger antigens with the number of residues greater than 400, while 3IRC is a smaller antigen with only 108 residues. Each of our training sets has a small number of antigens with sizes of >400 residues, and the predictions are thus expected to be poor for larger antigens. Therefore, the performance of ISPIPab appears to be size dependent and can be improved by including additional experimental data on both larger and smaller antigens, as they become available, to improve the predictive power across a wider range of antigen sizes.

## 4. Conclusion

In this work, we developed ISPIPab to antigen epitope prediction and show that our method outperforms several recent methods, including those designed specifically for B-cell epitope prediction, according to various statistical measures. Our method is partner-independent and predicts multiple epitopes on antigens based on the three-dimensional structures of unbound antigens. This is significant as knowledge of the cognate antibody or the three-dimensional structure of the antibody-antigen complex may not always be known.

Furthermore, multiple epitopes on a single antigen are identified using a hierarchical clustering methodology, where an optimal number of clusters are determined through dendrogram analysis. The results presented here demonstrate that clustering can improve prediction performance even in cases where only a single epitope is experimentally known. It is observed that often one predicted cluster of residues matches well with the experimentally identified epitope and the other cluster(s) may present a novel epitope where experimental data is not yet available. We also show that our methodology can accurately predict multiple distinct epitopes on a single antigen that agrees well with available experimental data. The availability of additional experimental data on very small and large antigens for training could further enhance the performance of ISPIPab.

## Acknowledgements

We acknowledge Mordechai Walder for reading the final manuscript as well as help with figures and Abraham Bodzin for DiscoTope calculations.

## Availability and Implementation

ISPIPab is implemented in Python and the code and sample data can be downloaded from https://github.com/mcarroll8/ISPIPab

## Funding

This work has been supported by the National Institutes of Health (NIH) grants GM136357 and AI141816.

## Conflict of Interest

none declared.

## Notes

https://github.com/mcarroll8/ISPIPab

### Competing Interest Statement

The authors have declared no competing interest.

