## Supplementary Information for "Computational Prediction of Multiple Antigen Epitopes"

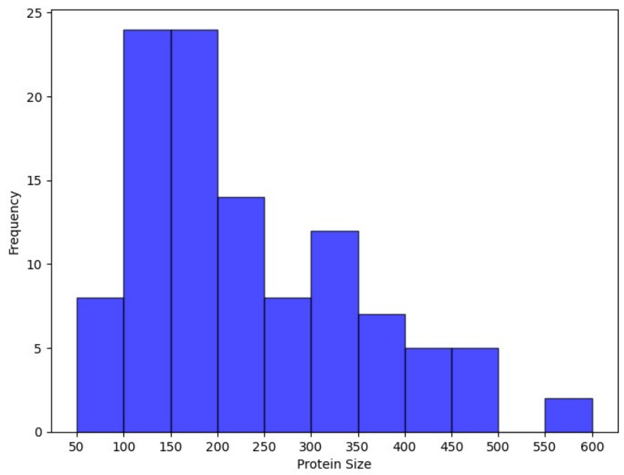


**Fig. S1. Size distribution of 109 unbound antigens in dataset B2.** Protein sizes ranging from 50 to 600 are shown. There are two proteins larger than 700 with the largest being 1277.

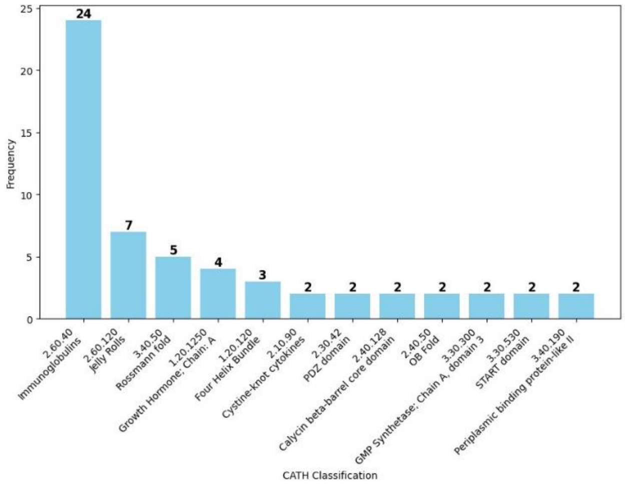


**Fig. S2**. **CATH classification (Orengo, Michie et al. 1997) of antigens in dataset B2**. Only those families represented by at least two antigens are included in the figure.

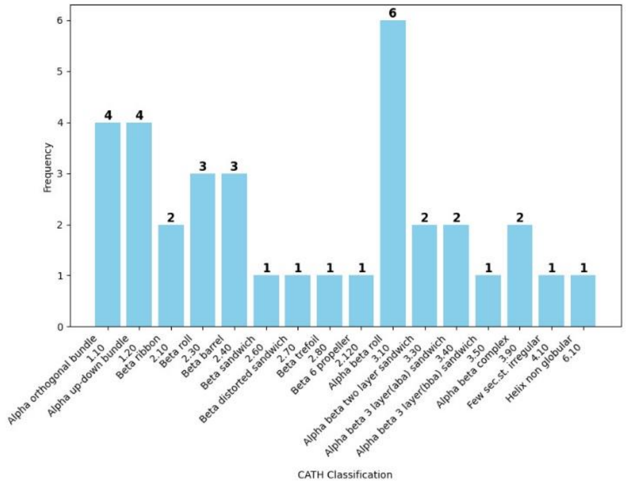


**Fig. S3. CATH Classification at the architecture level for the 35 antigens belonging to different topologies.**

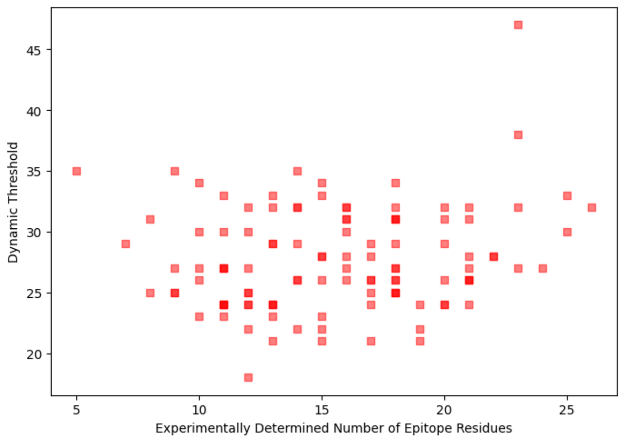


**Fig. S4. Comparison of the number of epitope residues predicted using the dynamic threshold with the annotated (experimentally determined number of) residues in unbound dataset B2.**

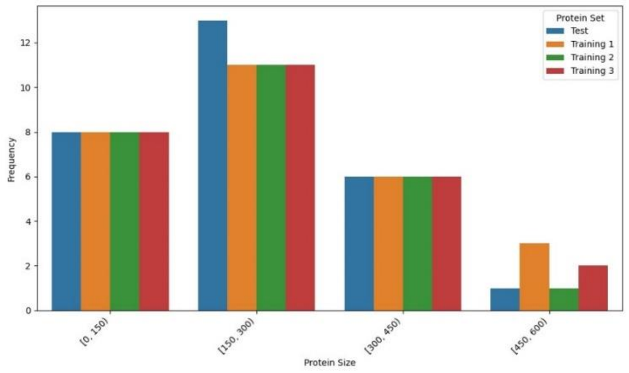


**Fig. S5. Size distribution of the antigens in each of the training and test sets**.
